# Acquired secondary HER2 mutations enhance HER2/MAPK signaling and promote resistance to HER2 kinase inhibition in HER2-mutant breast cancer

**DOI:** 10.1101/2022.09.23.509246

**Authors:** Arnaldo Marin, Abdullah Al Mamun, Hiroaki Akamatsu, Dan Ye, Dhivya R. Sudhan, Benjamin P. Brown, Lisa Eli, Katherine Marcelain, Jens Meiler, Carlos L. Arteaga, Ariella B. Hanker

## Abstract

*HER2* mutations drive the growth of a subset of breast cancers and are targeted with HER2 tyrosine kinase inhibitors (TKIs) such as neratinib. However, acquired resistance is common and limits the durability of clinical responses. Most *HER2*-mutant breast cancers progressing on neratinib-based therapy acquire secondary mutations in *HER2*. Apart from the *HER2^T798I^* gatekeeper mutation, whether these secondary *HER2* mutations are causal to neratinib resistance is not known. We show herein that secondary acquired *HER2^T862A^* and *HER2^L755S^* mutations promote resistance to HER2 TKIs via enhanced HER2 activation and impaired neratinib binding. While cells expressing each acquired *HER2* mutation alone were sensitive to neratinib, expression of acquired double mutations enhanced HER2 signaling and reduced neratinib sensitivity in 2D and 3D assays. Computational structural modeling suggested that secondary HER2 mutations stabilize the HER2 active state and reduce neratinib binding affinity. Cells expressing double HER2 mutations exhibited resistance to most HER2 TKIs but retained sensitivity to mobocertinib and poziotinib. Double-mutant cells showed enhanced MEK/ERK signaling which was blocked by combined inhibition of HER2 and MEK, providing a potential treatment strategy to overcome resistance to HER2 TKIs in *HER2*-mutant breast cancer.

## Introduction

Activating mutations in *HER2 (ERBB2)* are oncogenic drivers in ∼5% of metastatic breast cancers^1–4^. These mutations are primarily in the tyrosine kinase domain (KD) of the protein. In breast cancer, *HER2* mutations most often occur in the absence of *HER2* amplification, are more common in invasive lobular carcinoma (ILC)^5, 6^, and are associated with a worse prognosis^7–9^. Recurrent *HER2* mutations promote resistance to antiestrogen therapy in estrogen receptor-positive (ER+) breast cancer^10, 11^.

HER2 is a member of the ERBB receptor tyrosine kinase (RTK) family, which includes EGFR, HER3 (*ERBB3*), and HER4 (*ERBB4*). HER2 itself has no known ligand, but exists in an open conformation, enabling heterodimerization with ligand-bound HER family members^12^. Receptor dimerization enables transphosphorylation of tyrosine residues in the KD and subsequent signal transduction through the phosphoinositide-3-kinase (PI3K)/AKT/mTOR and RAS/RAF/ MEK/ERK signaling pathways^1^.

The irreversible pan-HER tyrosine kinase inhibitor (TKI) neratinib has demonstrated anti-tumor activity in pre-clinical models of *HER2-*mutant breast cancer^1, 13, 14^ as well as in clinical trials^3, 15, 16^. However, clinical responses to neratinib are short-lived and frequently associated with the acquisition of secondary mutations in *HER2*^15, 17^. While acquired ‘gatekeeper’ mutations are commonly known to promote acquired resistance to kinase inhibitors^18^ including neratinib in *HER2*-mutant cancers^2, 15^, the majority of acquired secondary mutations in HER2 are not in canonical gatekeeper residues (i.e., not T798I or L785F)^15^. Whether these non-gatekeeper secondary *HER2* mutations are <u>causal</u> to neratinib resistance is unknown. Thus, we hypothesized that secondary *HER2* mutations augment HER2 pathway activation and and/or alter HER2 TKI binding, thereby reducing sensitivity to HER2 TKIs.

## Results

### Acquired secondary HER2 mutations promote neratinib resistance

Two clinical trials in *HER2*-mutant breast cancers revealed high rates of acquired secondary *HER2* mutations in the circulating tumor DNA (ctDNA) following progression on neratinib-based therapy^15, 16^ (**Fig. S1A**). We chose to focus on tumors harboring HER2^L869R^ or HER2^L755S^, since multiple patients whose tumors harbored these primary mutations acquired secondary *HER2* mutations. Most of the secondary mutations we studied (shown in white in **Fig. S1**) have previously been demonstrated to activate HER2 signaling and promote breast cancer cell growth as single mutants^1, 2, 19^. In addition, single-mutant HER2^S310F^, HER2^D769Y^, HER2^D769H^, and HER2^L755S^ have been shown to be sensitive to neratinib *in vitro*^1, 19^, in contrast to the HER2^T798I^ gatekeeper mutation alone^2^.

We engineered MCF10A breast epithelial cells to stably express the *HER2^L869R^* or *HER2^L755S^* alone or together with acquired secondary mutations *in cis*^15^. We determined the neratinib IC_50_ for MCF10A HER2^L869R^, HER2^L755S^, and each double-mutant cell line. Expression of HER2^L869R/L755S^ and HER2^L869R/T862A^ shifted the neratinib IC_50_ (>15-fold) relative to HER2^L869R^ (**Figs. 1A, B**). Similarly, the HER2^L755S/T862A^ double mutant shifted the neratinib IC_50_ (>3.5-fold) relative to HER2^L755S^ (**Figs. 1C, D**). Therefore, we chose to focus on these three double mutants that promote neratinib resistance.

**Figure 1.**
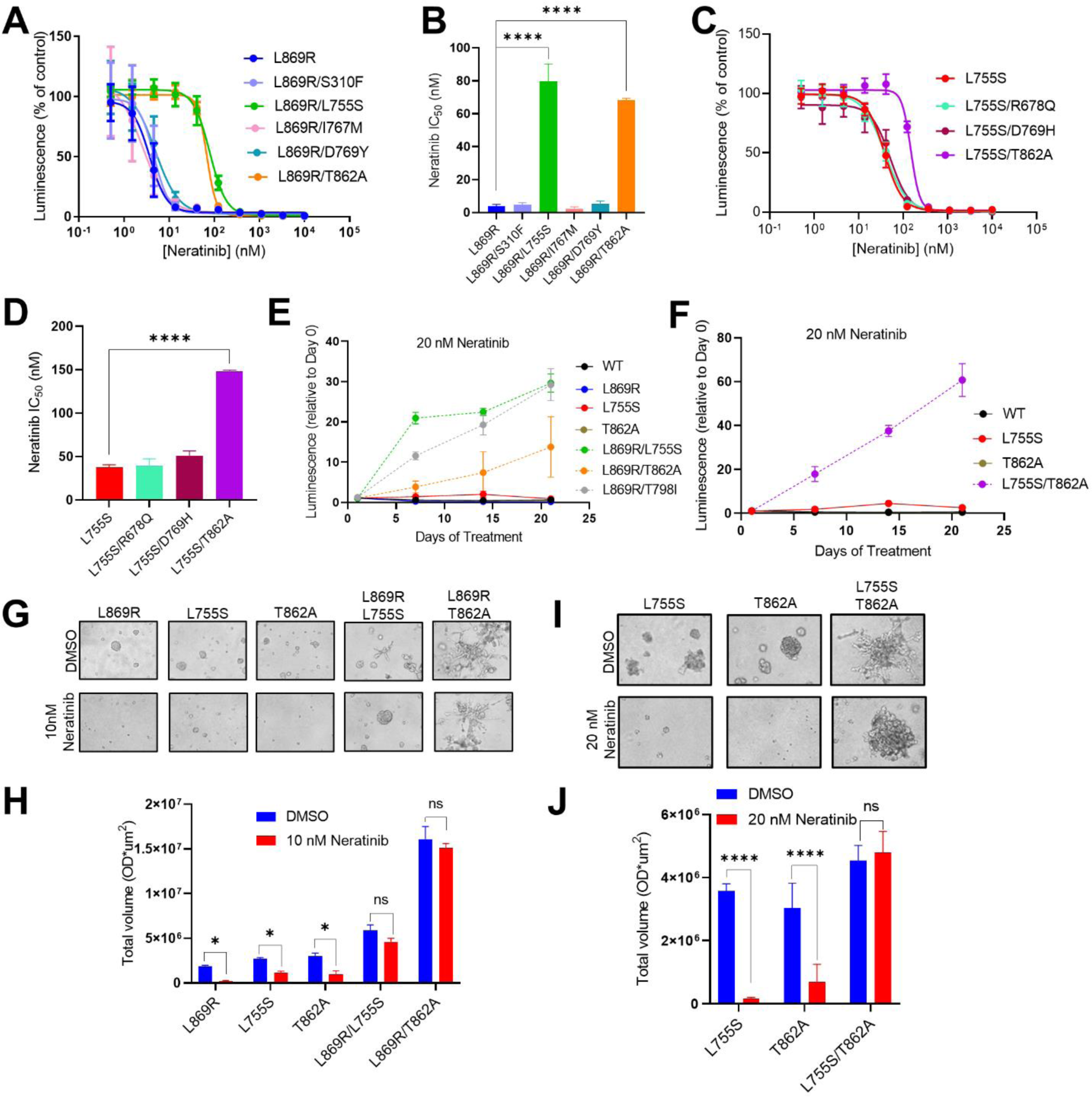
Acquired secondary HER2 mutations promote neratinib resistance. A-D) MCF10A cells stably expressing the indicated HER2 single- or double-mutants were grown in EGF/insulin-free medium + 1% CSS and treated with 11 concentrations of neratinib for 6 days. Cell viability was measured using CellTiter-Glo and IC_50_ values were calculated in GraphPad. Data represent the mean of three independent dose-response curves containing four replicates each. ****p<0.0001, ANOVA + Bonferroni. E) and F) MCF10A cells expressing the indicated HER2 constructs were grown as in the presence of 20 nM neratinib. Cell viability was measured every 7 days. Solid lines represent cells expressing HER2 single mutants; dotted lines represent HER2 double mutants. Data represent the mean ± SD (n=4). G) and I) MCF10A cells stably expressing the indicated constructs were grown in 3D Matrigel in EGF-free medium +1% CSS for 10 days in the presence of DMSO or neratinib 10 nM (G) or 20 nM (I). H) and J). The total volume of MTT-stained colonies per well was quantified using the GelCount instrument. Bars represent the mean ± SD (n=3). *p<0.05, ****p<0.0001, 2-way ANOVA + Bonferroni.

Growth in 2D of MCF10A cells expressing HER2^WT^ or each single mutant HER2^L869R^, HER2^L755S^, or HER2^T862A^) was inhibited by treatment with 20 nM neratinib (**Figs. 1E, F**). In contrast, cells expressing two mutations together (HER2^L869R/L755S^, HER2^L869R/T862A^, or HER2^L755S/T862A^) continued to expand during treatment with neratinib, like cells expressing the HER2 gatekeeper mutation (HER2^L869R/T798I^). Similarly, neratinib treatment suppressed the growth in 3D Matrigel of HER2 single mutants (HER2^L869R^, HER2^L755S^, or HER2^T862A^) but not double mutants (HER2^L869R/L755S^, HER2^L869R/T862A^, and HER2^L755S/T862A^) (**Figs. 1G-J**). Although the other double mutants examined _(HER2L869R/S310F, HER2L7869R/I767M, HER2L869R/D679Y, HER2L755S/R678Q, and HER2L755S/D769H)_ failed to significantly affect neratinib sensitivity in 2D, most displayed moderately reduced sensitivity to neratinib in 3D (**Fig. S2A, B**). These data suggest that simultaneous expression of two neratinib-sensitive HER2 mutations promotes neratinib resistance.

### Acquired secondary HER2 mutations enhance HER2 activation and signaling output

Expression of each HER2 single mutant, with the exception of HER2^S310F^ (in the extracellular domain instead of the KD), increased ligand-independent HER2 auto- phosphorylation relative to HER2^WT^; all single mutants enhanced phosphorylation of AKT, ERK, and S6, signaling molecules downstream of HER2 (**Figs. 2A, C**). Most of the double mutants further enhanced AKT, ERK, and S6 phosphorylation relative to HER2^L869R^ or HER2^L755S^ single mutants (**Figs. 2A-D**), suggesting that acquired secondary *HER2* mutations enhance HER2 signaling.

**Figure 2.**
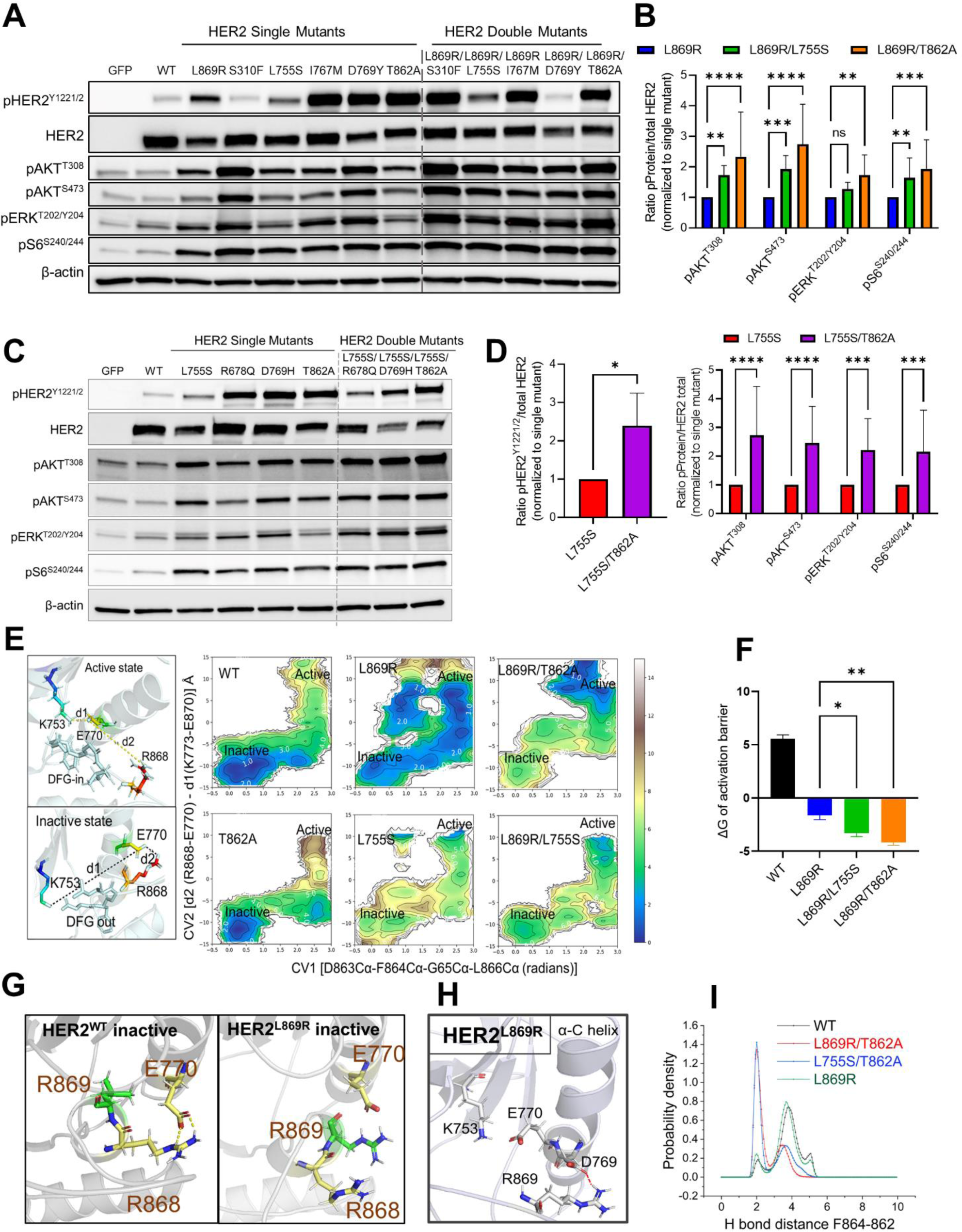
Acquired secondary HER2 mutations enhance HER2 activation. A) and C) Lysates from MCF10A cells expressing the indicated constructs were probed with the indicated antibodies. B) and D) Immunoblot bands from three independent experiments were quantified using ImageJ. Ratios relative to the ectopic expression of HER2 are shown to account for differences in HER2 expression. Ratios were normalized to immunoblots from cells expressing the HER2 single-mutant (HER2^L869R^ or HER2^L755S^). E) Free energy landscape (FEL) for HER2^WT^ and mutants. Two different collective variables (CV) representing the active and the inactive states are shown in the left panel. The right panel represents the FEL for the active to inactive state conformational transition. F) Quantification of the energy barrier (ΔG) for the conformational transition from active to inactive states. The barrier was calculated based on the absolute energy of the active and inactive states based on the equation: active state minimum energy – inactive state minimum energy. G) The **inactive state** structure of HER2^WT^ and L869R was obtained from MD simulation. H) The structural model of HER2^L869R^ in the **active state**. The interaction between R869 and D769, which plays a prominent role in the stabilization of the α-C helix, is shown. I) The probability density between F864-T/A862 distances for the indicated mutants was plotted by combining three 1μs MD replicates. *p<0.05, **p<0.01, ***p<0.001, ****p<0.0001, 2-way ANOVA + Bonferroni.

To understand how the double mutants augment HER2 activation, we applied computational structural analysis of HER2^WT^ and HER2-mutant models. The structure and the location of the primary and secondary mutations are shown in **Figure S2C**. Conventional and enhanced molecular dynamics (MD) simulations were employed using the monomeric model of the HER2 KD. To explore the effect of each mutation on the conformational changes from the HER2 active to the inactive state, we applied umbrella sampling (US), a form of enhanced MD^20^. This was used to calculate the two-dimensional free energy landscape (FEL) of the WT, single, and double mutants. The detailed methodology for the US, including the collective variables (CV), is discussed in the Methods section. The US simulations were based on the trajectories obtained from the steered molecular dynamics (SMD) simulations from the HER2 active to an inactive state and vice-versa. The FEL was generated with two different collective variables (CV1 and CV2), where CV1 consists of the dihedral angle between the D863-F864-G865-L866 (DFG-motif) and CV2 is the difference in distances between K753-E770 and R868-E770 (left panel, **Fig. 2E**). These two CVs, the conformation of DFG-motif and canonical salt bridge in the α-C helix are important parameters to differentiate the active and inactive state conformations in ErbB receptors^21–23^.

FEL modeling demonstrated that the inactive state of HER2^WT^ is more stable compared to the active state (difference of ∼5 kcal/mol), as evidenced by the low energy minima in the inactive state (CV1 0.25, CV2 -13) (**Figs. 2E, F**). Similar results were reported using free energy studies for EGFR and HER2^21, 22, 24^. The FEL of HER2^T862A^ was similar to that of HER2^WT^, with distinct inactive state minima (CV1 0.25, CV2 -12) which represent the inactive state stabilization compared to the active state. In contrast, the inactive state of HER2^L869R^ is destabilized, potentially because R869 engages in an unfavorable interaction with the two-turn helix of the activation loop, thus preventing the formation of the R868-E770 salt bridge and favoring the active state (**Figs. 2G, H**). In addition, in the HER2^L869R^ active state, residues R869 and D769 form favorable interactions that preserve the canonical salt bridge K753-E770 in the α-C helix which may stabilize the active state (**Fig. 2G**). The active state of HER2^L869R/T862A^ and HER2^L869R/L755S^ was *further* stabilized compared to L869R alone (**Fig. 2E**). This effect appears to be driven by destabilization of the inactive state in the double mutants. The presence of A862 with R869 causes additional favorable interactions which stabilize the activation loop and thus the active state (**Fig S2D**). Using cMD simulations for HER2^L869R/T862A^, we found that a backbone hydrogen bond between A862 and F864 that facilitates the extended conformations of the activation loop (**Fig. S2D**). The probability distribution between residues T/A862-F864 was notably higher for HER2^L869R/T862A^ aggregated over all simulation replicates (**Fig. 2I**), suggesting H-bond formation is more dominant due to that mutation.

Next, we determined the ATP binding affinities for single and double mutants. We conducted 1μs cMD simulations for the ATP-bound state. We found that HER2^L869R/L755S^ and HER2^L755S/T862A^ double mutants are predicted to exhibit increased ATP-hinge interactions (**Figs. S2E-G**). ATP-hinge interactions are important for ATP binding and a strong interaction is indicative of higher activity. We further calculated the ATP binding affinity for HER2 WT and mutants. Based on molecular mechanics Poisson Boltzmann surface area (MMPBSA) simulations, the ATP binding affinity for HER2^L755S/T862A^ was predicted to be enhanced compared to HER2^L755S^ (Fig. S2G).

### Acquired secondary HER2 mutations are incompletely blocked by neratinib

Next, we examined the effect of neratinib on HER2 signaling induced by the double mutants. Fifty (50) nM neratinib was sufficient to completely block phosphorylation of HER2, HER3, AKT, ERK, and S6 in MCF10A cells expressing HER2 single mutations (HER2^L869R^, HER2^L755S^, or HER2^T862A^). In contrast, treatment with 50-100 nM neratinib, which exceeds the C_max_ in patients^25^, failed to completely block phosphorylation of the same molecules in cells expressing double mutants (HER2^L869R/L755S^, HER2^L869R/T862A^, or HER2^L755S/T862A^; **Fig. 3A,B**). Similarly, even high concentrations of neratinib (100 nM) failed to effectively block the growth in 3D Matrigel of cells expressing the double mutants (**Fig. S3A-D**).

**Figure 3.**
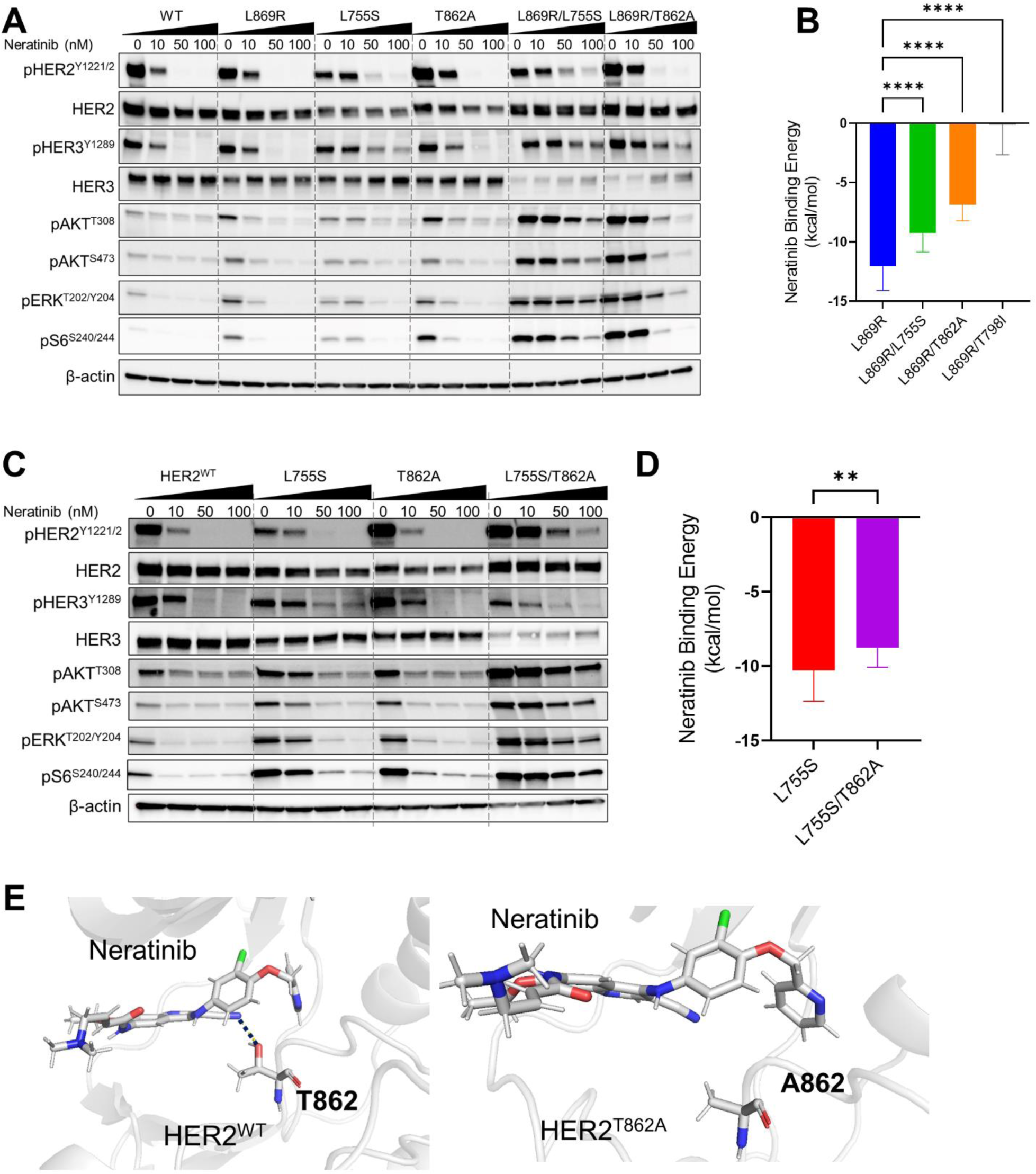
Acquired secondary HER2^L755S^ and HER2^T862A^ mutants are not blocked by neratinib. A) and C) MCF10A cells stably expressing HER2^WT^ or the indicated mutants were treated with DMSO or the indicated concentrations of neratinib for 4h. Lysates were probed with the indicated antibodies. B) and D) Molecular mechanics Poisson Boltzman surface area (MMPBSA) and simulation was performed to estimate the binding affinity of neratinib to HER2 single- or double-mutants. E) The neratinib interaction with Thr862 of HER2 (blue dotted line) is abrogated in the HER2^T862A^ mutant (A862). The structures were taken from MD snapshot after 1 μs of MD simulations. **p<0.01, ****p<0.0001, 2-way ANOVA + Bonferroni.

To determine whether acquired secondary HER2 mutations affect neratinib binding, we applied cMD simulations. We first examined the structural perturbation induced by HER2^T862A^. Our simulation suggested that the T862 residue in HER2^WT^, HER2^L869R^, and HER2^L755S^ single mutants forms a van der Waals interaction with neratinib (**Fig. S3E**). We also observed that HER2 residues S783 and T862 form a van der Waals interaction with the nitrile group of neratinib (**Fig. S3E**). The T862A mutation impacts the sidechain orientation of S783 as well as the interaction of the S783 residue with neratinib (**Fig. S3F**). To ensure that the T862-neratinib interaction is an energetic minimum, we applied cMD simulations with different initial conformations of the T862 residue in HER2^WT^. We initially modified the sidechain orientation of T862 to a position that is unfavorable for interaction with the nitrile group of neratinib and used this structure as input for cMD (**Fig. S3G**). However, after 100 ns of cMD simulation, the sidechain of T862 changes orientation such that it favorably interacts with neratinib suggesting that the secondary HER2^T862A^ mutation negatively impacts neratinib binding due to the loss of the T862-neratinib interaction.

We next applied MMPBSA simulations to calculate neratinib binding^26^. These simulations suggested that HER2^T862A^ destabilizes the interactions with neratinib (**Figs. 3C, D**). We further asked whether this neratinib-T862 interaction is the only limiting factor for drug resistance. To understand the contributions of the HER2-T862A interaction, we calculated a per residue energy breakdown for residues that are potentially important for neratinib binding (**Figs. S3H, I**). Energy decomposition analysis suggested that S783, T862, M801, D863 favorably interact with neratinib and are responsible for the higher neratinib binding affinities of HER2^WT^, HER2^L869R^, and HER2^L755S^. The HER2^T862A^ mutation impacted the overall binding energy and the interaction between S783 and neratinib (**Figs. 3C-E** and **S3E**). In contrast, the reduced neratinib binding affinity of HER2^L869R/L755S^ is the result of structural changes, as evidenced by stabilization of the active state along with destabilization of the neratinib-D863 interaction (**Figs. 2E** and **S3H, I**). In both HER2^L869R/T862A^ and HER2^L869R/L755S^, the K753-E770 salt bridge is stabilized due to the favorable interactions of the L755S and R869 in the active state (**Fig. S3J**). Our per residue analysis suggests that this unfavorably affects the D863 interactions with neratinib and hence reduces the binding affinity (**Fig. S3I**). In addition, the greater affinity of HER2^L755S/T862A^ for ATP discussed above (**Fig. S2I**) would negatively affect the binding of neratinib, an ATP-competitive inhibitor. Together, our results suggest that acquired secondary HER2^T862A^ and HER2^L755S^ mutations induce neratinib resistance by both enhancing HER2 activation and by negatively impacting neratinib binding.

### Acquired secondary HER2 mutations reduce sensitivity to most HER2 TKIs

There are currently around ten pan-HER or HER2-selective TKIs in clinical trials for *HER2-*mutant tumors (clinicaltrials.gov). We screened ten concentrations of nine structurally distinct TKIs in MCF10A cells expressing HER2^L869R^ or HER2^L755S^ primary _mutations and HER2L869R/L755S, HER2L869R/T862A, HER2L869R/T798I, and HER2L755S/T862A_ double mutants. Compared to the single mutants, cells expressing the HER2^L869R/L755S^, HER2^L869R/T862A^, and HER2^L755S/T862A^ double mutants showed reduced sensitivity to the majority of TKIs tested (**Figs. 4A** and **S4A**). Cells expressing the HER2^L869R/T798I^ gatekeeper mutation, which directly impedes drug binding^2^, did not show reduced sensitivity to smaller TKIs including such as afatinib, tarloxitinib, and poziotinib. Therefore, we hypothesized that the reduced sensitivity of the HER2^L869R/L755S^, HER2^L869R/T862A^, and HER2^L755S/T862A^ double mutants may in part be due to enhanced HER2 activation, which is not seen in the HER2^L869R/T798I^ mutant^2^ (**Figs. 4B** and **S4B-E**).

**Figure 4.**
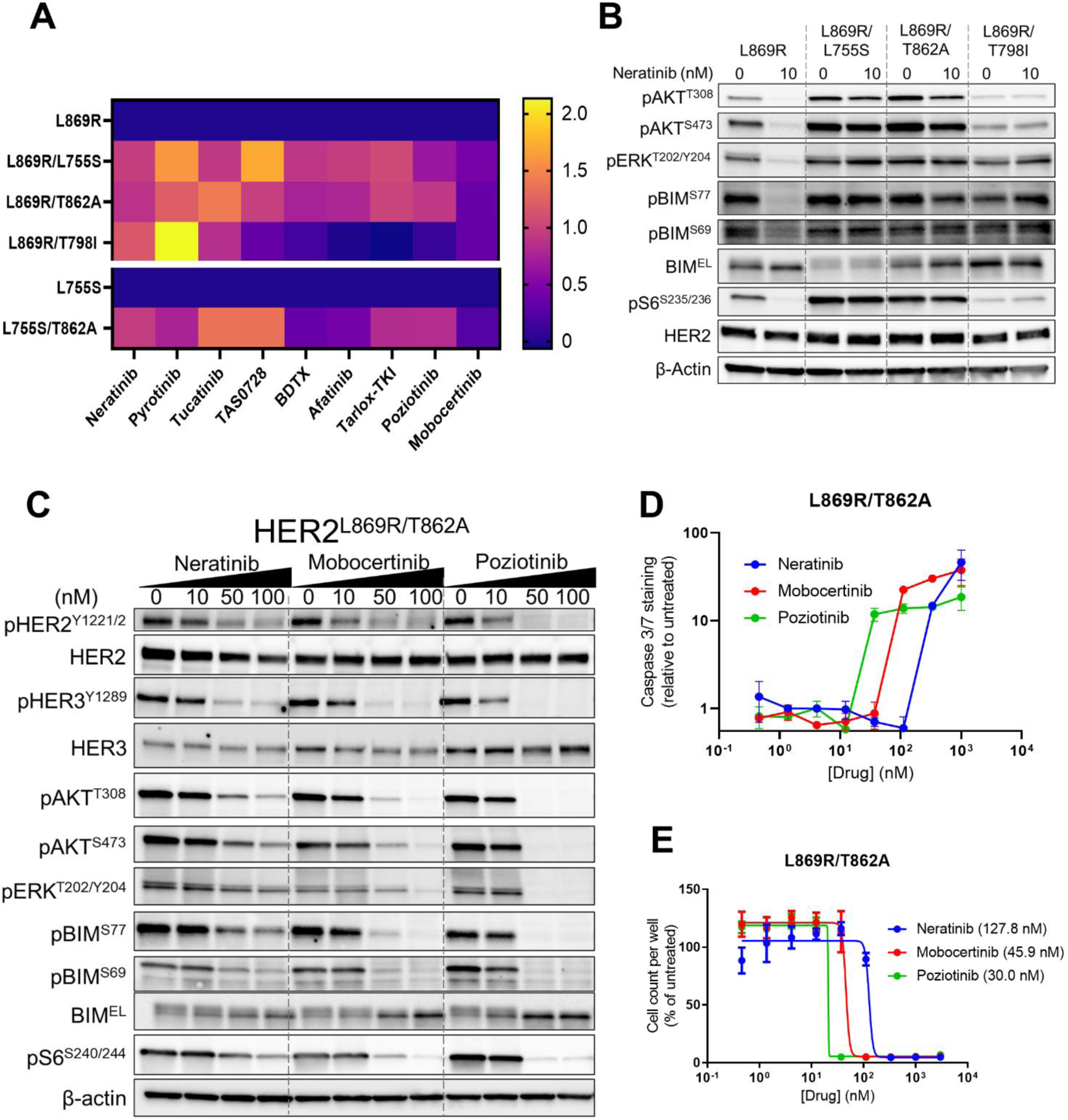
Acquired secondary HER2 mutations reduce sensitivity to most HER2 TKIs, but mobocertinib and poziotinib. A) MCF10A cells stably expressing the indicated HER2 mutants were treated with 10 increasing concentrations of each HER2 tyrosine inhibitor (TKI) for 6 days. A heat map of the Log (IC_50_) normalized to the primary L869R or L755S mutation from duplicate CellTiter-Glo viability assays is shown. B) MCF10A cells stably expressing HER2^L869R^ single- or double-mutants were treated with DMSO or 10 nM neratinib for 4h. Lysates were probed with the indicated antibodies. C) MCF10A HER2^L869R/T862A^ cells were treated with the indicated concentrations of each inhibitor for 4 h. D) MCF10A HER2^L869R/T862A^ cells were treated with the indicated concentrations of each inhibitor and stained with Caspase3/7 Green fluorescent reagent. Fluorescence was quantified after 3 days of treatment. Data points represent the Caspase3/7-stained area/number of total cells per well, relative to untreated cells (n=3). E) Cells were treated as in (D). The total number of cells per well after 6 days of treatment was quantified by Incucyte Cell-by-Cell Analysis (n=3).

Cells expressing HER2 double mutants retained sensitivity to clinically achievable doses of mobocertinib^27^ and poziotinib^28, 29^ (**Figs. 4A** and **S4A**). Both TKIs blocked phosphorylation of AKT, ERK, S6, and BIM more potently than neratinib in cells expressing double HER2 mutants (**Figs. 4C** and **S4F-H**). The pro-apoptotic BIM protein is a substrate of ERK (**Fig. S4I**) and inhibition of phospho-BIM has been suggested as a potential biomarker of sensitivity to HER2 inhibition in HER2-amplified breast cancer^30–32^. ERK-mediated BIM phosphorylation results in ubiquitin-mediated BIM degradation, and inhibition of ERK increases expression of the pro-apoptotic extra-long BIM (BIM_EL_)^33, 34^. Indeed, treatment with lower doses of poziotinib or mobocertinib than neratinib induced apoptosis (as measured by caspase 3/7 staining) in HER2 double-mutant cells (**Figs. 4D** and **S4J**). This was associated with enhanced sensitivity to poziotinib and mobocertinib (**Figs. 4E** and **S4K**).

Poziotinib is a second generation EGFR/HER2 covalent inhibitor^35, 36^ and mobocertinib is third generation EGFR/HER2 covalent inhibitor^37, 38^. We applied cMD simulations followed by MMPBSA to calculate the binding affinity of mobocertinib and poziotinib. Based on the binding affinity prediction, the binding affinity of neratinib and mobocertinib to the HER2 kinase should follow similar topology (**Fig S5A-B**). However, this appeared to be different for poziotinib, where we observed similar binding affinities for the single and double mutants (**Fig. S5A,C,D**), consistent with the activity of poziotinib against cells with double HER2 mutations. Analysis of per residue interactions showed that T798, D863 and T862 residues plays crucial role in poziotinib binding (**Fig. S5C,D**). In contrast to neratinib, the T862 residue is not predicted to directly interact with poziotinib, offering a rationale for more favorable binding of the T862A-containing double mutants.

In contrast to neratinib, both mobocertinib and poziotinib displayed more favorable binding energies to the HER2^L869R/T798I^ gatekeeper mutant (**Fig. S5A**). The favorable binding of mobocertinib with HER2^L869R/T798I^ is attributed to the strong interactions of T862 with the mobocertinib linker group (**Fig. S5E**). Similarly, poziotinib shows strong interactions with residues D863 and M801 in HER2^L869R/T798I^ (**Fig. S5F**). These results suggest that mobocertinib and poziotinib may be more effective therapies for breast cancers with double HER2 mutations.

### HER2 double-mutant cells are dependent on the MEK/ERK pathway

We next asked which downstream pathways are required for the growth of HER2 double-mutant cells. Since HER2 double-mutant cells exhibited enhanced phosphorylation of ERK, AKT, and S6, we tested inhibitors of MEK (selumetinib or trametinib), PI3Kα (alpelisib), AKT (ipatasertib or capivasertib), and mTOR1 (everolimus), alone or in combination with neratinib. HER2 single- and double-mutant cells were more sensitive to MEK inhibitors (MEKi) than PI3Ki, AKTi, or mTORC1i, alone or in combination with neratinib (**Figs. 5A, B** and **S6A-C**). We previously showed that cells expressing concurrent HER2/HER3 mutations are highly sensitive to the combination of neratinib and alpelisib^22^, but the HER2 double-mutant cells were relatively insensitive to this combination (**Fig. S6D**). Mechanistically, only MEKi blocked ERK-induced phospho-BIM and induced apoptosis in double-mutant cells (**Figs. 5C-E** and **S6E-H**), while both MEKi and mTORC1i reduced cell proliferation (**Fig. S6I**).

**Figure 5.**
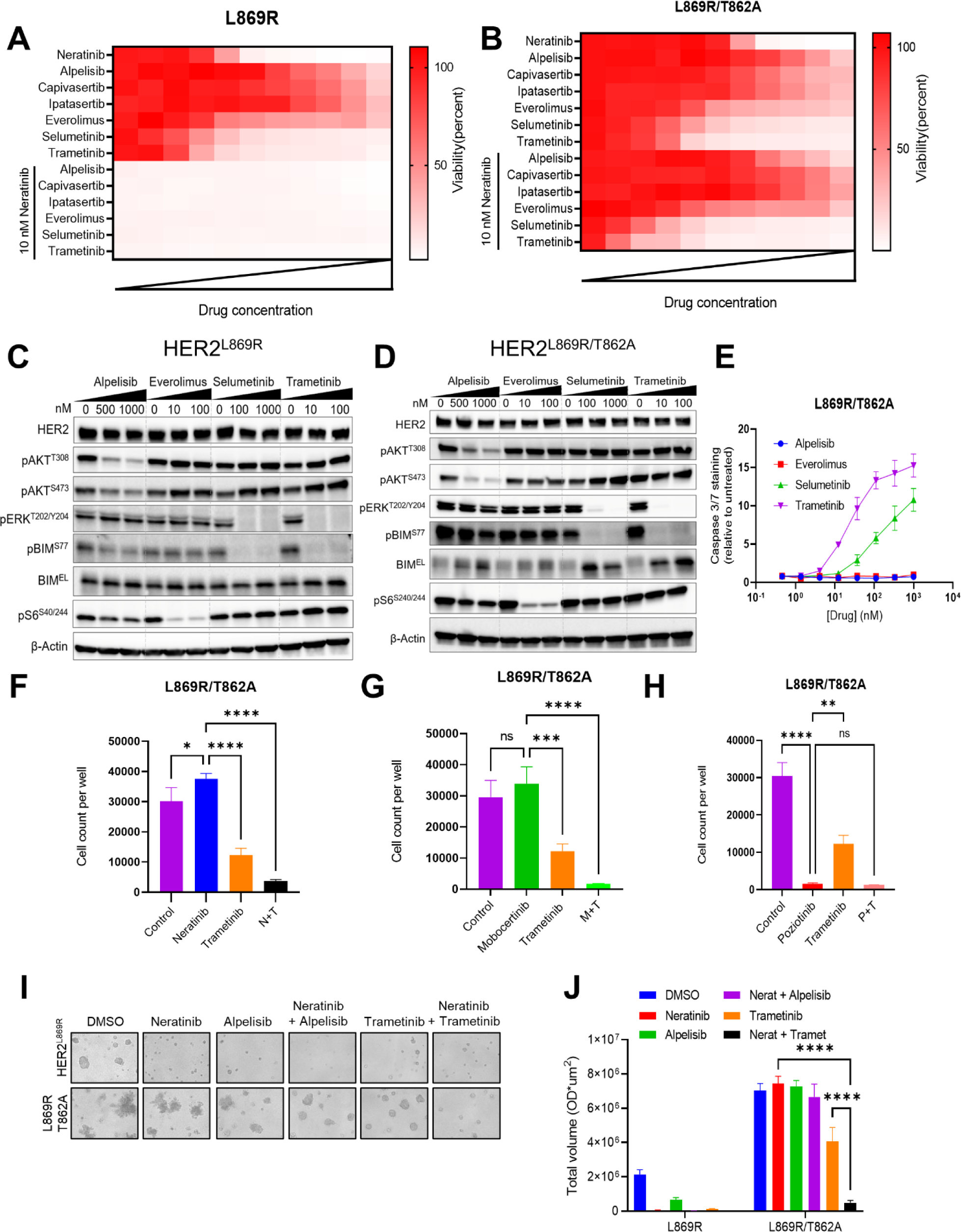
HER2 double-mutants are sensitive to MEK inhibition. A-B) MCF10A cells stably expressing the indicated HER2 mutants were treated with increasing concentrations (3-fold) of each drug, up to 10 μM. For combination assays, all cells were treated with 10 nM neratinib and increasing concentrations of the second drug. Cell viability was measured by the CellTiter-Glo assay. Data represent the mean of quadruplicate wells. C-D) MCF10A cells stably expressing D) HER2^L869R^ and E) HER2^L869R/T862A^ were treated with the indicated inhibitors for 4 h. Lysates were probed with the indicated antibodies. E) MCF10A HER^L869R/T862A^ were treated with the indicated concentrations of inhibitors and stained with Caspase3/7 Green fluorescent reagent. Fluorescence was quantified after 3 days of treatment. Data points represent the Caspase3/7-stained area/number of cells per well, relative to untreated cells (n=3). F–H) MCF10A HER^L869R/T862A^ cells were treated with concentrations of each HER2 TKI (Neratinib 37 nM, Mobocertinib 37 nM or Poziotinib 37 nM) ± trametinib (12 nM). The number of cells per well after 6 days of treatment was quantified by Incucyte. Data represent the mean ± SD (n=3). I) MCF10A cells stably expressing HER2^L869R^ single- and double-mutants were grown in 3D Matrigel in EGF-free medium +1% CSS for 10 days and treated with DMSO, 10nM Neratinib, 1uM Alpelisib, and 10nM Trametinib alone or in combination. J) The total volume of MTT-stained colonies per well was quantified using the GelCount instrument. Data represent the mean ± SD (n=3). *p<0.05, **p<0.01, ***p<0.001, ****p<0.0001, 2-way ANOVA + Bonferroni.

Finally, we examined the impact of MEK inhibition in combination with neratinib mobocertinib, or poziotinib. The combination of trametinib with neratinib or mobocertinib more effectively suppressed HER2 double-mutant cell proliferation than either drug alone (**Fig. 5F-H**). Similarly, the combination of neratinib and a MEKi more effectively suppressed growth of HER2 double-mutant cells in 3D Matrigel than either drug alone, whereas the combination of neratinib and alpelisib failed to suppress growth (**Fig. 5I, J**). Together, our data suggest that that drug-resistant secondary mutations in *HER2* amplify RAS/RAF/MEK/ERK signaling, and inhibition of this pathway may delay or reverse resistance to HER2 TKIs, suggesting a testable therapeutic strategy for this subtype of breast cancers.

## Discussion

Resistance to TKIs is frequently driven by secondary gatekeeper mutations in the target oncogene which occlude drug binding while maintaining oncogene gain-of-function and dependence^39–41^. HER2 is no exception—the HER2^T798I^ secondary gatekeeper mutation has been found in several *HER2*-mutant breast cancer patients following progression on TKI-based therapy^2, 15–17^. However, unlike frequent gatekeeper mutations in other common oncogenes (EGFR^T790M^; KIT^T670I^), HER2 gatekeeper mutations are found in a minority of patients following progression. Instead, secondary HER2 mutations in sites that are known to activate HER2 activity are quite frequent in HER2-mutant breast cancer patients progressing on neratinib^15, 17^. Here, we systematically tested acquired secondary HER2 mutations found in patients with clinically resistant tumors and discovered that most double mutants enhanced HER2 signaling and oncogenesis while reducing sensitivity to neratinib (**Figs. S2A, B and 2A, C**). In particular, acquired mutations in HER2^T862A^ and HER2^L755S^ strongly promoted resistance to neratinib (**Fig. 1**). These mutants alone are sensitive to neratinib (**Figs. 1E-J**), suggesting that secondary acquired mutations in HER2^T862A^ and HER2L^755S^ impair sensitivity to the ATP-competitive inhibitor in part through an increased threshold of ATP binding and kinase activity. Such a mechanism was previously hypothesized by Hyman and colleagues but was not tested experimentally^15^. We found that HER2^L869R/T862A^, HER2^L869R/L755S^, and HER2^L755S/T862A^ double-mutant cells were relatively resistant to at least seven other HER2 TKIs (**Fig. 4A**). These double mutants also exhibited reduced neratinib binding affinity (**Figs. 3B, D**). Thus, we conclude that <u>both</u> reduced neratinib binding and enhanced ATP binding and HER2 activation play a role in neratinib resistance. This mechanism is distinct from the HER2 gatekeeper mutant, which solely impedes neratinib binding but does not enhance HER2 activation^2^ (**Figs. S4B-D and 4B**) and remains sensitive to other HER2 TKIs (**Fig. 4A**). This mechanism is also distinct from concurrent HER2/HER3 mutations, which reduce neratinib binding affinity while enhancing HER3/PI3K/AKT activation and dependence on HER2 and PI3K signaling^22^, whereas HER2 double mutants are insensitive to this combination but strongly inhibited by combined blockade of HER2 and MEK (**Figs. 5 and S6**).

There are several examples of increased activation of oncogenic kinases mediating resistance to kinase inhibitors. Perhaps the best-known examples are the acquired amplification of the mutant allele of BRAF^V600E^ in melanomas following progression on BRAF inhibitors^42, 43^ and EGFR amplification following treatment with EGFR inhibitors^44^. Similarly, kinase domain duplications of BRAF have been shown to mediate BRAF inhibitor resistance^45^. Interestingly, secondary drug-resistant mutations in BRAF are quite rare. In these cases, amplification of the target is thought to limit drug efficacy by exceeding its inhibition capacity^46^. Similarly, we show that higher concentrations of the HER2 TKIs, often exceeding their cMax, were needed block signaling and growth induced by the double mutants. To our knowledge, this study is the first report of secondary kinase *missense mutations* that reduce drug sensitivity primarily via enhanced target activation rather than reduced drug binding.

The HER2^L755S^ single mutation has been associated with a lesser response to neratinib therapy in clinical studies^17^. In line with this, we and others previously showed that MCF10A cells expressing HER2^L755S^ are less sensitive to neratinib relative to cells expressing other HER2 oncogenic missense mutations^19, 22^. In addition, L755 or its homolog in other kinases has been identified as a conserved site whose substitution results in resistance to ATP-competitive kinase inhibitors^47^. Our structural modeling suggests that in HER2^L755S^, the N-terminal region of the α-C helix is stabilized, which preserves the inward conformation of the helix, resulting in a higher ATP binding affinity and steric repulsion with neratinib (**Figs. S2E-J and S3J**). However, cells expressing HER2^L869R/L755S^ and HER2^L755S/T862A^ are clearly more resistant to HER2 TKIs relative to cells expressing HER2^L755S^ alone, suggesting that both an increase in HER2 kinase activity and a modest reduction in neratinib binding affinity contribute to drug resistance.

The EGFR^T854A^ mutation is homologous to HER2^T862A^, which we showed abolishes the interaction between HER2 S783 and neratinib, leading to a moderate decrease in neratinib binding affinity (**Figs. S3E-F**). Notably, EGFR^T854A^ has previously been associated with resistance to EGFR inhibitors^48–50^. HER2^T862A^ is also resistant to BI-1622, a novel HER2 inhibitor shown to directly interact with S783^51^. Based on computational structural studies for HER2 mutations, we have shown that neratinib directly interacts with T862 and S783 (**Figs. S3E-F**). Substitutions of the threonine with alanine impede the neratinib-T862 interaction and destabilizes the S783 orientation which, in turn, affects the neratinib-S783 interaction. We that speculate BI-1622 (not available for this study) may not directly interact with T862. Instead, we posit that the T862A mutation destabilizes the interaction between HER2 S783 and the HER2 TKI, as shown for neratinib (**Figs. S3E-I**). Higher concentrations of neratinib, which are not clinically achievable, were needed to block the growth of cells with these double mutations. In contrast, we found that cells expressing these double mutants were sensitive to clinically relevant doses of poziotinib and mobocertinib. Therefore, poziotinib or mobocertinib may lead to more durable responses compared to neratinib for *HER2-*mutant breast cancers and/or may overcome acquired resistance and, as such, warrant clinical investigation in *HER2-*mutant breast cancers.

HER2 double mutations have also been found in neratinib-naïve patients^52, 53^ and have been associated with intrinsic drug resistance in the SUMMIT trial^3, 15^. In the Project GENIE database^54^, ∼9% of primary HER2-mutant breast cancers harbor double HER2 mutations, including 5 patients with L755S and 2 patients with T862A as one of the two HER2 mutations in the tumor. We posit that many of these HER2 double-mutant tumors will also be less sensitive to HER2 TKIs due to enhanced HER2 activation. We would also anticipate that with increased use of HER2 TKIs in patients with *HER2*-mutant cancers and use of next-gen sequencing of metastases that progress after an initial clinical response, the frequency of acquired drug-resistant secondary mutations should increase.

We showed that HER2 single- and double-mutant cells were both strongly dependent on the MAPK pathway, as MEK inhibition by either selumetinib or trametinib was more effective at reducing cell viability compared to inhibition of PI3K, AKT, or mTOR (**Figs. 5A-B**). This is consistent with previous studies showing that MCF7 cells expressing mutant HER2 activate a MAPK gene signature^11^. In contrast, HER2^WT^-amplified breast cancer cells are very dependent on the PI3K pathway and sensitive to PI3K/AKT inhibitors^55–57^. In agreement, we previously showed that *HER2* mutations alone did not strongly induce HER3/PI3K activation^22^. Smith et al. recently reported that *HER2*- amplified tumors harboring MAPK pathway alterations, including *HER2* mutations, displayed a switch from PI3K to MAPK pathway dependence^58^, consistent with our findings in *HER2* double-mutant cells.

In summary, we found that secondary *HER2* mutations acquired during neratinib progression in patients with breast cancer enhance HER2 activation and signaling to PI3K/AKT and MEK/ERK pathways and reduce sensitivity to HER2 TKIs. These acquired mutations result in a higher threshold of HER2 tyrosine kinase activation, and inhibition of this signaling output requires a higher concentration of HER2 TKIs. The HER2 double-mutants were exquisitely sensitive to the combination of HER2 and MEK inhibitors. The combination of neratinib and trametinib demonstrated strong efficacy in patient-derived HER2-amplified xenografts^59^ and is currently in clinical testing for HER2-mutated or amplified tumors (NCT03065387). Our results also suggest that other HER2 TKIs like poziotinib or mobocertinib may allow for a high enough dose to disable the activation of HER2 double-mutants and reduce or delay on-target resistance. Development of mutant-selective HER2 inhibitors may allow for a wider therapeutic index than drugs that block WT HER2/EGFR, allowing for more complete inhibition of HER2 single- and double-mutants and more durable clinical responses.

## Materials and Methods

### Cell lines

MCF10A cells were purchased from ATCC. Cell lines were authenticated by ATCC prior to purchase by the tandem short repeat method. 293FT cells were purchased from Invitrogen.

293FT cells were maintained in DMEM supplemented with 10% FBS and 1x antibiotic-antimycotic. MCF10A cells were maintained in complete MCF10A medium (DMEM/F12 supplemented with 5% horse serum, 20 ng/ml EGF, 10 mg/ml insulin, 0.5 mg/ml hydrocortisone, 0.1 mg/ml cholera toxin and 1X antibiotic/antimycotic). For all experiments, MCF10A cells were grown in EGF-free DMEM/F12 supplemented with 1% charcoal/dextran-stripped serum (CSS), 0.5mg/ml hydrocortisone, 0.1 mg/ml cholera toxin and 1X Antibiotic-Antimycotic. Cell lines were routinely evaluated for Mycoplasma contamination. All experiments were completed less than 2 months after establishing stable cell lines or early passage thawing of such cells.

### Plasmids

The Gateway Cloning system (Thermo Fisher Scientific) was used to generate the pLX302-HER2 plasmids. The pDONR-223 vector encoding HER2^WT^ was subjected to site-directed mutagenesis (Genewiz) to generate HER2 single or double mutants *in cis*. HER2^WT^ and the mutant plasmids were recombined into the lentiviral expression vector pLX-302 containing a C-terminal V5 epitope tag and puromycin resistance marker.

### Lentiviral infections

Lentiviral supernatant was produced in early passage 293FT cells by transfection with packaging plasmids psPAX2 and pMD2.G, along with the appropriate plasmid pLX302- HER2. The next day, target cells were spin-infected with viral supernatant in the presence of 8 mg/ml polybrene. Two days later, target cells were selected with 2 μg/ml puromycin for at least 4 days. Stable cell lines were maintained in media containing puromycin.

### Western blot analysis

MCF10A cells were grown in EGF-free media + 1% CSS for at least 24 hours prior to lysing. Adherent cells were washed with ice-cold PBS and lysed with NP40 buffer (Sigma) supplemented with 1X protease inhibitor (Roche) and phosphatase inhibitor cocktails (Roche). Lysates were centrifuged at 13,500 rpm for 5 min. Protein concentrations in supernatants were quantified using the BCA protein assay kit (Pierce). 20-40 mg of total protein was fractionated on 4-12% bis-tris gradient gels (BioRad) and transferred to nitrocellulose membranes. Membranes were blocked with 5% nonfat dry milk/TBST at room temperature for 1 h, followed by overnight incubation with antibodies at 4°C in 5%BSA/TBST. All antibodies were purchased from Cell Signaling: pHER2^Y1221/2^ (#2243; 1:500), HER2 (#2242; 1:1000), pEGFR^Y1173^ (#4467; 1:500), EGFR (#4267; 1:1000), pHER3^Y1289^ (#4791; 1:500), HER3 (#12708; 1:1000), pAKT^S473^ (#9271; 1:500), pAKT^T308^ (#13038; 1:500), p-S6^S240/244^ (#5364; 1:1000), pERK^T202/Y204^ (#9101; 1:1000), BIM (#2939;1:1000), pBIM^S69^ (#4585;1:500) pBIM^S77^ (#12433; 1:500) and β-actin (#4970; 1:1000). Membranes were sliced horizontally to probe with multiple antibodies. The antibodies for pAKT^S473^, pERK ^T202/Y204^, and pS6^S240/244^ were combined during primary incubation. Nitrocellulose membranes were washed and incubated with HRP-conjugated rabbit or mouse secondary antibodies for 1 hour at room temperature. Protein bands were detected with an enhanced chemiluminescence substrate (Perkin Elmer) using the ChemiDoc Imaging System (Bio-Rad). Immunoblots were quantified using ImageJ.

### Cell viability and apoptosis assays

MCF10A cells were grown in EGF-free media + 1% CSS. Cell viability was determined using the CellTiter-Glo luminescent assay (Promega) according to the manufacturer’s instructions. Single-cell suspensions were generated by filtering trypsinized cells through a 40 mm cell strainer (Fisher Scientific). 500-1000 cells per well were seeded into white clear-bottom 96-well plates in quadruplicate. Cells were treated with 10 concentrations of inhibitor or vehicle control at a final volume of 150 μl per well. After 6 days of treatment, 25 μl of CellTiter-Glo reagent was added to each well. The Caspase-Glo 3/7 Assay (Promega) was performed according to the manufacturer’s instructions. Plates were shaken for 15 min and bioluminescence was determined using the GloMax Discover microplate reader (Promega). Blank-corrected bioluminescence values were normalized to DMSO-treated wells; normalized values were plotted on GraphPad Prism 9 using a nonlinear regression fit with variable slope (four parameters). IC_50_ values were calculated by GraphPad Prism.

Cell proliferation and apoptosis were measured using the Incucyte S3 Live-Cell Analysis System. One thousand cells per well were seeded in clear 96-well plates (Corning) in triplicate. The next day, cells were labeled with Caspase-3/7 Dye (for apoptosis) and Nuclight Rapid Red Dye (for live-cell nuclear labeling) (Incucyte) and up to 11 concentrations of drugs were added for dose-response assays. Apoptosis was quantified after 3 days of treatment by calculating the Caspase 3/7-stained area divided by the number of total cells per well. Cell proliferation was quantified after 6 days of treatment.

### Three-dimensional morphogenesis assay

Cells were seeded in growth factor-reduced Matrigel (Corning #356230) in EGF-free media + 1% CSS in 48-well plates, following published protocols^25^. Inhibitors were added to the medium at the time of cell seeding. Fresh media and inhibitors were replenished every 3 days. After 7-10 days, colonies were stained with 5 mg/ml MTT for 20 min. Plates were scanned and colonies measuring ≥ 100 μm were quantified using GelCount software (Oxford Optronix). Colonies were photographed using a Leica DMi1 inverted microscope.

### Quantification and statistical analysis

Statistical analysis was performed with GraphPad Prism 9. For analyses involving multiple comparisons, one-way or two-way (for grouped bar graphs) ANOVA with Bonferroni’s posthoc test was used. Bar graphs show mean ± S.D. Drug combination indices were calculated using the Chou-Talalay test^26^.

### Computational methods

#### Model preparation

HER2 is a single-pass transmembrane protein with an intracellular kinase domain and is well known for adopting a wide range of conformations such as active, inactive, and src-like inactive states based on the structural flexibility of the kinase domain^60–65^. These three different states are typically dictated by the conformation of the activation loop, the DFG motif, and the stability of the α-c helix. The distinct feature of an active state is an extended conformation of the activation loop, an inward conformation of the α-c helix to facilitate the canonical salt bridge K753-E770 formation, and inward sidechain conformation for the ASP in the DFG motif (Fig. S7A). In the inactive state, the alpha-c helix shifts to an outward conformation which breaks the salt bridge formation between the K753-E770, and the activation loop adopts a double helix conformation (Fig. S7B). A HER2 active state crystal structure has not yet been solved. To model this active state conformation of HER2, we initially build the active state of HER2 using comparative modeling in Rosetta with PDB structures 4riw and 3pp0 as a template^66, 67^. Two rounds of comparative modeling were conducted where the best-scored models from the first iteration were used as a template for the second round of comparative modeling^68^. 10000 models were generated and clustered using distance-based rmsd as a similarity metric (Fig. S7C). The best-scored model from each cluster was further refined using 100 ns cMD simulation. The best-performing structures were then further relaxed using Rosetta fast relax and used for subsequent modeling of HER2 active state.

To prepare the inactive state model, we further used Rosetta comparative modeling protocol using crystal structures 2gs7 and 3rcd as a template^61, 65, 69^. The final model for subsequent molecular simulation was further refined using the similar protocol described above for the HER2 active state (Fig. S7D). To introduce the single and the double mutation in HER2, Rosetta ddg monomer was used. The structures were relaxed, and the mutations were introduced using ddg monomer which predicts the structural flexibility of the mutant with respect to the WT analog^70^. The best-scored model from ddg monomer were further relaxed using Rosetta fast relax and used for the initial input preparation for MD simulation.

#### Inhibitor docking

Three different covalent inhibitors, neratinib, mobocertinib, and poziotinib were used with HER2 active and inactive state models. For neratinib, a second-generation covalent inhibitor, the initial pose for the inhibitor binding was obtained from the crystal structure (PDB ID- 3W2Q)^71^. The initial structure of the neratinib was first superimposed with HER2 WT and the mutants. The complexed structure was then relaxed using Rosetta and the best-scored model from the Rosetta relax was used as the initial input for the MD simulation. To dock mobocertinib with HER2, we initially used the osimertinib structure to superimpose with HER2 models. Mobocertinib is a third-generation covalent inhibitor and to date, there is no crystal structure in complex with HER2/EGFR is available. We initially used osimertinib structure to build the mobocertinib model and the structure was optimized using the quantum mechanical method. The structure was optimized using density functional theory (B3LYP/6-311g(d))^72, 73^ followed by Rosetta ligand docking with HER2 based on the known binding pose of the osimertinib (PDB ID: 6Z4B)^74^ and the best-scored model from the docking used for the MD simulation. Poziotinib, a second-generation covalent inhibitor was similarly docked with HER2 using Rosetta, and the best-scored model from Rosetta ligand docking was used for the MD simulation.

#### MD simulation

The initial structure obtained from Rosetta was subsequently used for the molecular dynamics (MD) simulation with the FF14SB force field using the Amber16.0 software suite^75, 76^. Three different model systems, protein-inhibitor complex, apo-protein, and protein-ATP complex in different HER2 mutants were used for the MD simulation. For protein-inhibitor complexes, three different inhibitors that were used in the simulation are, neratinib, mobocertinib, and poziotinib. The initial structure of the inhibitors were first optimized using the B3LYP/6-311g(d) level of theory in Gaussian 16 ^77^. The optimized structure was used for the parameterization of the protein-inhibitor complex using the GAFF force field.

The model systems were solvated with a 10 Å TIP3P water box and Na^+^/Cl^-^ counter ions were added for the neutralization. The initial models were first minimized with constraints for 10000 steps with steepest descent algorithms followed by 5000 steepest conjugate gradient method. The model system was then minimized without constraints for 15000 steepest-descent followed by 5000 conjugate gradient method. The minimized structure was then heated from 50 K to 100 K in 100 ps in a constant number of molecules, constant volume, and constant temperature (NVT) ensemble using the 1fs time step. Subsequently, the system was heated from 100 K to 300 K for 1ns using the 1fs time step. Following the heating, the systems were equilibrated/production runs were conducted for 1 μs using an NPT ensemble. For each case, three separate simulations were performed with a random seed for velocities. For heating and production run using NPT ensemble, periodic boundary conditions were applied, and the long-range electrostatic interactions were calculated using particle-mesh Ewald (PME) algorithms with a 10Å cut-off for non-bonding interactions. Shake algorithms were used to constraint the hydrogen bond stretching. Production runs were performed in a 4fs time scale using hydrogen mass repartitioning in amber. Trajectories were analyzed and processed using the built-in program in amber CPPTRAJ^78^. VMD was used for the analysis of each trajectory and pymol to visualize the structure of different snapshots from MD simulation^79, 80^.

### Enhanced sampling

#### Steered molecular dynamics (SMD)

While conventional MD is quite good for the conformational study of biomolecules, however for many biomolecules with the rough, energetic landscape, it is often difficult to overcome the high energy barrier to sample different local minima^20^. Long molecular simulation is often trapped in a rough energy landscape which is nonfunctional for studying conformational transition at a reasonable computational time. Enhanced molecular dynamics simulations such as steered molecular dynamics (SMD) or umbrella sampling (US) can overcome this barrier by introducing a harmonic potential^81^. SMD applies an external potential and pulls a specific configuration at a constant velocity towards a reference configuration. The time-dependent potential using a force constant k is, 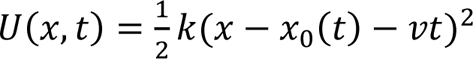. Here, x is the reference configuration, *x*_0_(*t*) is the initial configuration at time t and k is the spring constant. Thus, if the spring constant is large enough, after a certain time the initial configuration will reach the final configuration. Here in this method, we applied four different SMD simulations for the active to inactive state conformational transition. To sample the overall conformational space of the HER2, we applied SMD with different force constants. For each mutant four SMD simulations were performed, two from active to inactive and vice versa with two different force constants. The initial setup of the SMD is similar as described in the MD setup. Following the heating and equilibration with NVT and NPT ensemble, the constant velocity SMD was performed for 10 ns using 2000 kcal/mol/A force constant from the active state towards inactive state conformations and vice versa. The collective variable that was used for the SMD is Cα rmsd of the total system. Additionally, we performed SMD for 10 ns using a 500 kcal force constant from active to an inactive state transition.

#### Umbrella sampling (US)

Umbrella sampling is an enhanced sampling technique where an additional potential is applied to the system by dividing total simulation into different fragments each of which represents a window^82^. For each of these windows, a simple harmonic potential V_i_ is added which is dependent on the force constant k of the system. The harmonic potential 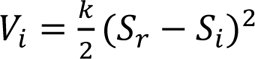, here *V_i_* is the harmonic potential, *k* is the force constant, *S_i_* is the reference distances/angles and *S_r_* is the actual distance/angle of the collective variables (CV). Two different collective variables (CV1 and CV2) were used to generate the FEL from active to inactive state transition. The choice of the collective variable is based on the active and inactive state conformation.

For CV1, D863-F864-G865-L866 dihedral angle was used as a collective variable, since this motif adopts two different conformations in the active and inactive state (Fig. S7A-B). In the case of an active state, the side chain of D863 adopts an inward conformation, which plays a critical role in ATP binding. In an inactive state, the motif changes toward outward conformation. These two conformations are the distinctive active and inactive state features and hence a crucial coordinate in FEL generation. For CV2, two salt bridge distances were used as the coordinate. In the active state conformation, K753-E770 forms a salt bridge and stabilizes the αC-helix. The K753-E770 salt bridge is completely broken in the inactive state, similarly, a new salt bridge R868-E770 formation takes place (Fig. 2E). The differences in distances between these two salt bridges represent the active and inactive state position and are used as CV2 in constructing FEL. To construct the FEL, 300 windows were used with two different force constants (k), for CV1 and CV2 such as 20 kcal/mol and 2kcal/mol respectively. The selection of windows is based on the SMD simulations as described earlier. Initially, we applied SMD to sample the overall conformational landscape of HER2 from an active to an inactive state and vice versa. Each of the SMD simulations was divided into different windows and used as a starting point for umbrella sampling simulation. For each mutant 300 windows were used to generate the free energy landscape (FEL). Each of the simulations was used for 5ns production run at a constant temperature of 310K and the constant pressure of 1 atm. Hydrogen mass repartitioning was used to control the integration time step of 4 fs. The temperature was controlled using Langevin dynamics and the shake algorithm was used to constraint the bond involving hydrogen. Each of these simulations was then used for the potential of mean force (PMF) construction using the WHAM scheme as implemented in the reference^83^. To construct the PMF, the first 1ns simulation were discarded and the rest of the 4 ns were used. To calculate the standard deviation, each of the 5 ns simulations, were fragmented by 1ns timescale and WHAM was implemented to calculate the PMF^83^. The delta G of the simulation is calculated based on the absolute energy differences of the active and the inactive state. The positive (+) delta G indicates the inactive state stabilization, and the negative (-) delta G refers to the active state stabilization.

#### Binding affinity prediction

Molecular mechanics Poisson-Boltzmann surface area (MMPBSA) was applied to calculate the binding affinity of the ATP, neratinib, mobocertinib, and poziotinib^26^. Prior to this, the inhibitor was first docked with the WT and the mutant protein followed by cMD simulations for each of the mutants. We randomly resampled 1000 structures for each of the mutant-drug complex and applied MMPBSA simulation using MMPBSA.py module in Amber16. For MMPBSA simulations, the ionic strength was set to .15 mM, and 50 trajectories were used for each MMPBSA simulation. Overall, for each protein-drug complex, 20 MMPBSA simulations were conducted, and statistical analyses were performed for the average binding affinity prediction.

## Acknowledgments

We received the following financial support: NCI R01CA224899 (C.L. Arteaga and A.B. Hanker), NCI Breast SPORE P50 CA098131 (C.L. Arteaga and A.B. Hanker, UTSW Simmons Cancer Center P30 CA142543, CPRIT RR170061 (C.L. Arteaga), Susan G. Komen Breast Cancer Foundation SAB1800010 (C.L. Arteaga), and grants from the Breast Cancer Research Foundation (C.L. Arteaga and A.B. Hanker). Jens Meiler is supported by a Humboldt Professorship of the Alexander von Humboldt Foundation.

## Supplementary Figures

**Supplementary Figure 1.**
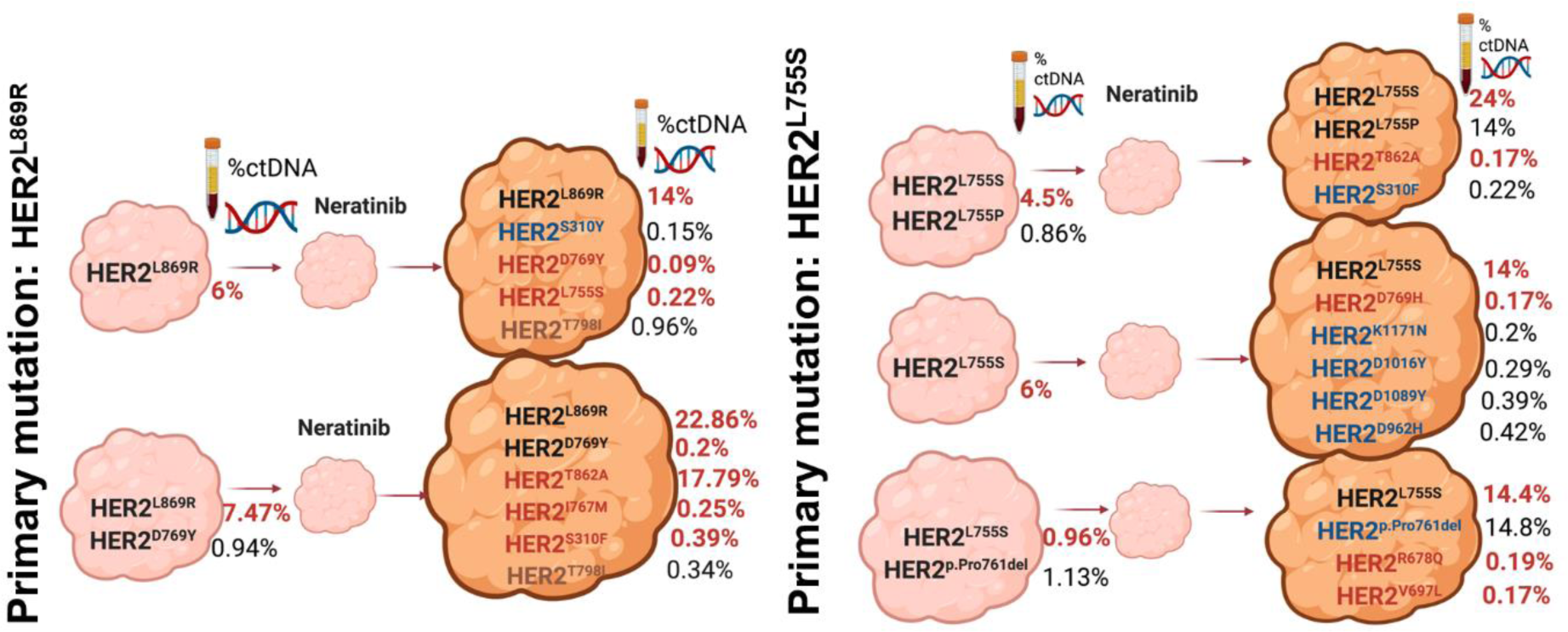
Acquired secondary HER2 mutations following neratinib progression in HER2-mutant breast cancer patients. Acquired secondary *HER2* mutations detected following neratinib-containing regimens in clinical trials of ER+/*HER2*-mutant breast cancer (compiled from the works of Smyth et al., *Cancer Disc.* 2020 Feb;10(2):198-213 and Ma et al., *Clin Cancer Res.* 2017 Oct 1;23(19):5687-5695). The allele frequencies of *HER2* mutations in circulating tumor DNA (ctDNA) pre-treatment and after progression are shown. Acquired secondary mutations that were investigated are shown in red. The gatekeeper mutation is shown in brown.

**Supplementary Figure 2.**
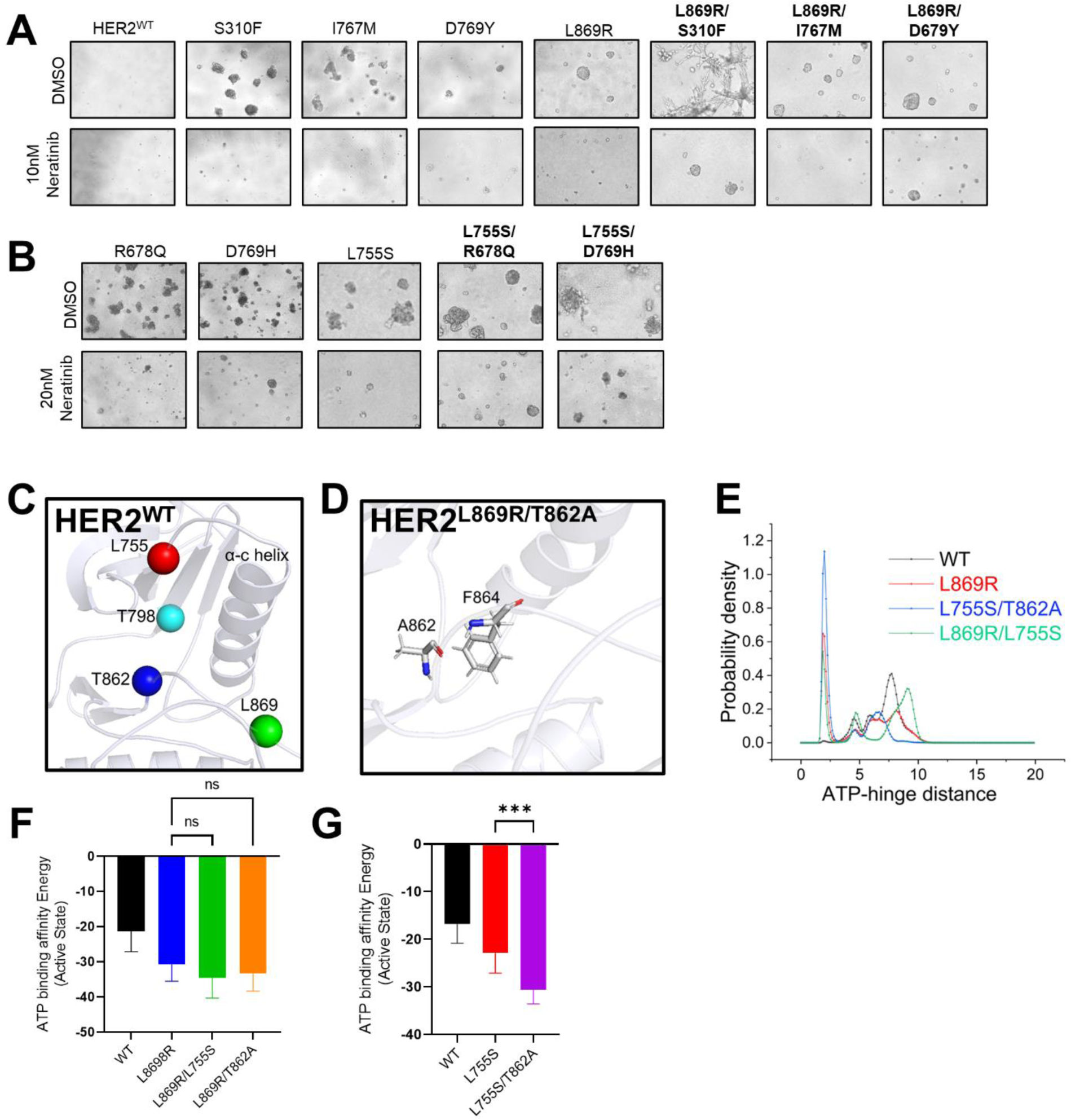
Acquired secondary HER2 mutations enhance HER2 activation. A) MCF10A cells stably expressing the indicated HER2^L869R^ single- and double-mutants were grown in 3D Matrigel for 10 days and treated with DMSO or 10 nM neratinib. B) MCF10A cells stably expressing the indicated HER2^L755S^ single- and double-mutants were grown in 3D Matrigel for 10 days and treated with DMSO or 20 nM neratinib. C) The active state of HER2 and the location of the KD mutations in HER2 D) The interaction between F864-A862 in HER2^L869R/T862A^. E) The probability density of the ATP-hinge region for different HER2 mutations. F-G) The ATP binding affinity for HER2 single- and double-mutants was calculated by MMPBSA simulations. The trajectories for MMPBSA simulation was obtained from cMD simulation coupled with hidden Markov model to identify the most probable long-lived states. ***p<0.001, 2-way ANOVA + Bonferroni

**Supplementary Figure 3.**
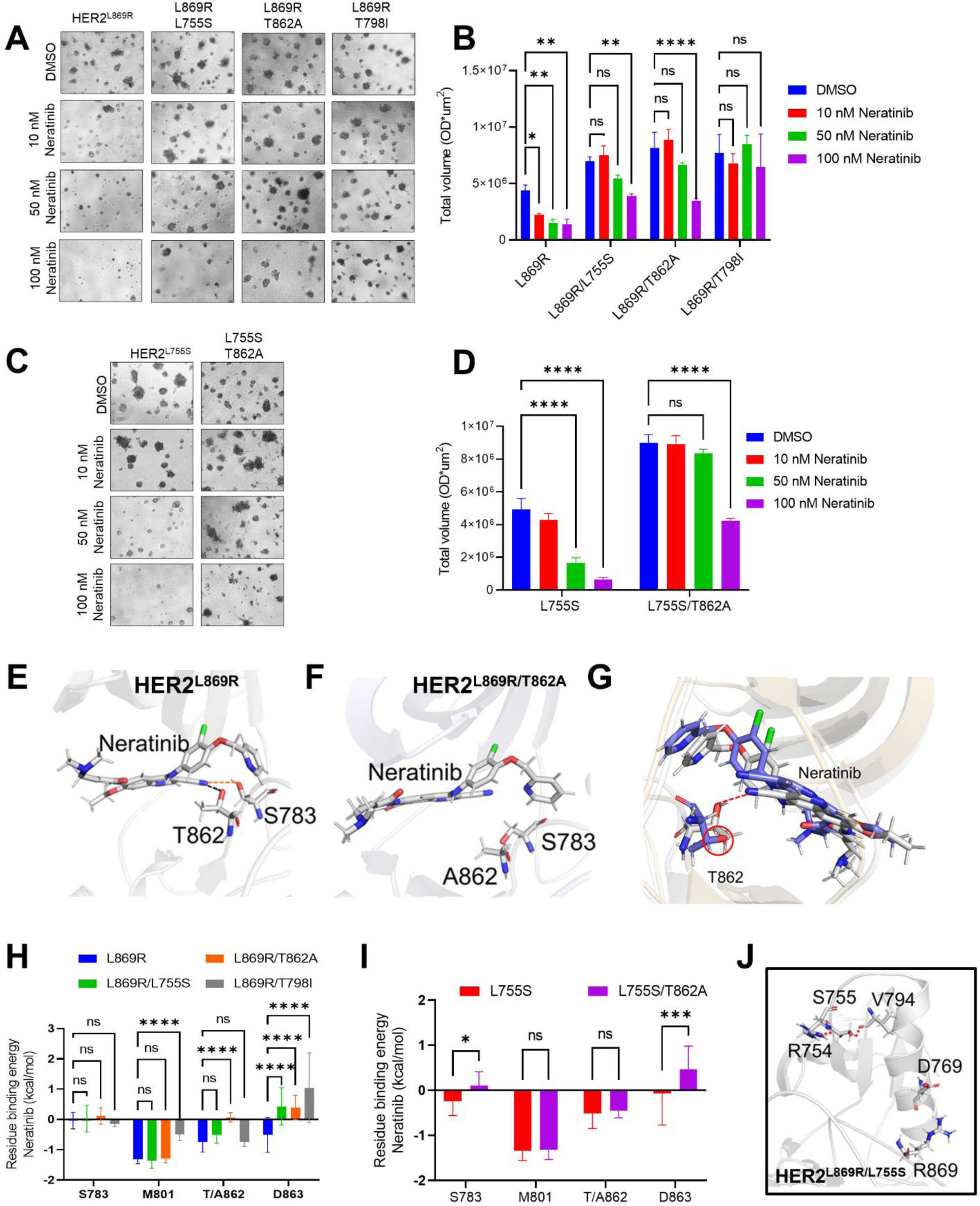
Acquired secondary HER2^L755S^ and HER2^T862A^ mutations are incompletely blocked by neratinib. A) and C) MCF10A cells stably expressing the indicated constructs were grown in 3D Matrigel and treated with DMSO or neratinib at the indicated concentrations. B) and D) The total volume of MTT-stained colonies per well was quantified using the GelCount instrument. Bars represent the mean ± SD (n=3). E) The structural model of HER2^L869R^ bound to neratinib. The snapshot is obtained after 1μs MD simulation. F) The structural model of HER2^L869R/T862A^, showing destabilization of the interaction between neratinib and S783. G) Two different conformations of HER2^WT^ are superimposed and the interaction of T862 with neratinib is highlighted. The green model represents the structure where the T862 orientation was changed to prevent the interaction and used as an input for MD simulation. The structural model in gray is after 100 ns MD simulation, where T862 returns in the conformation, which is favorable for making the interaction. H-I) Energy decomposition analysis for HER2 residues S783, M801, T/A862, and D863, to show the effect of each mutation on the contributions of these residues to neratinib binding. The energy decomposition analysis was performed by MMPBSA simulation. J) The structural model of HER2^L869R/L755S^. The interaction of R754-S755-V794 in the N-terminal region of the α-C helix is shown. *p<0.05, **p<0.01, ***p<0.001, 2-way ANOVA + Bonferroni.

**Supplementary Figure 4.**
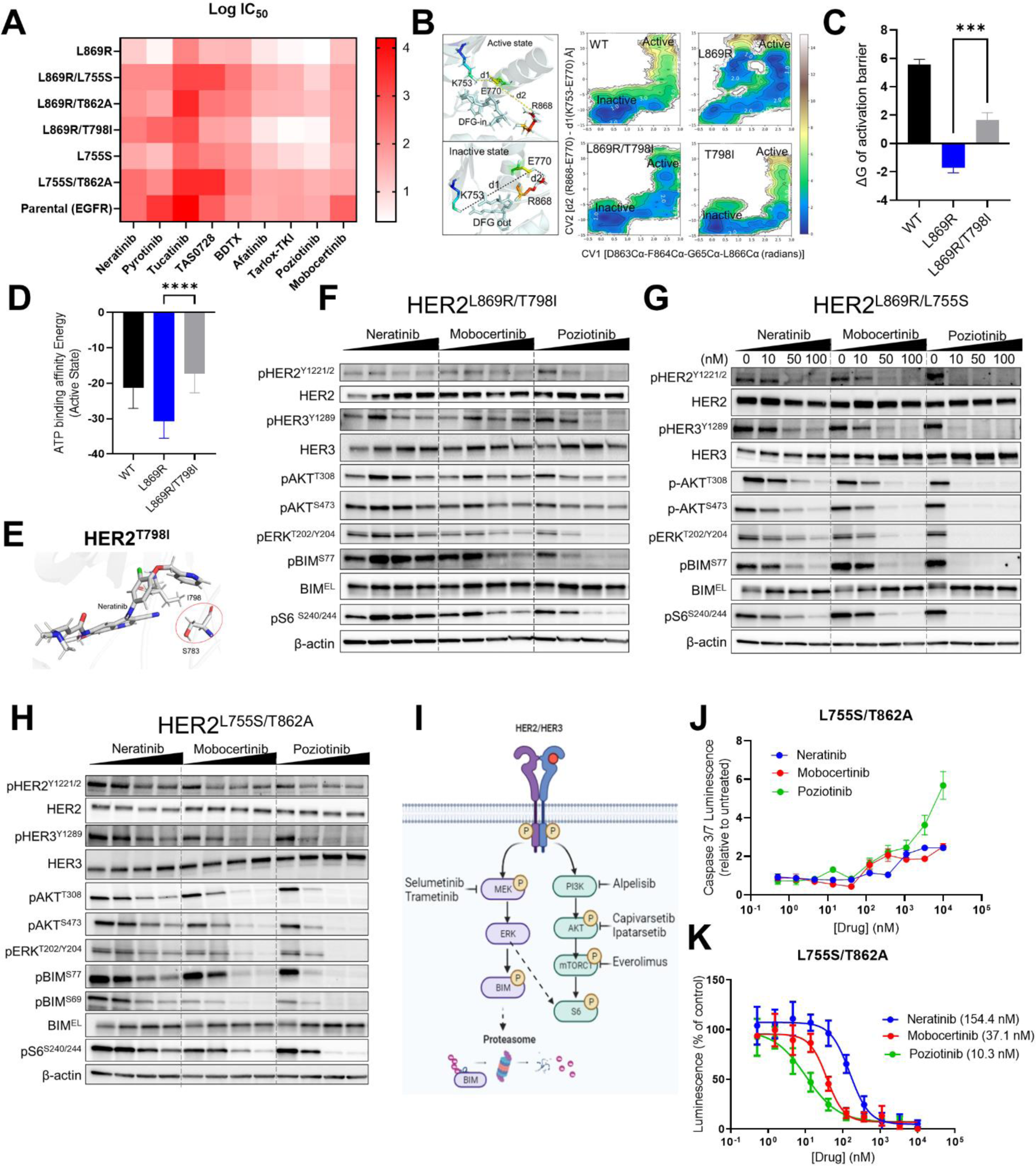
Acquired secondary HER2 mutants retain sensitivity to mobocertinib and poziotinib. A) MCF10A cells stably expressing the indicated HER2 mutants were treated with 10 increasing concentrations of each HER2 tyrosine inhibitor (TKI) in EGF/insulin-free media + 1% CSS for 6 days. MCF10A EGFR-overexpressing cells were grown in the presence of EGF. A heat map of the Log(IC_50_) from duplicate CellTiter-Glo viability assays is shown. B) FEL for HER2^WT^ and the indicated mutants. The collective variable representative of the active and inactive state is shown in the left panel. C) The conformational energy barrier for the active to inactive state transition is shown. D) The ATP binding affinity of HER2^WT^ and the indicated mutants was predicted by MMPBSA simulations. E) The structural model of HER2^L869R/T798I^. The sidechain orientation of S783 is disrupted by I798. F-H) MCF10A cells stably expressing the indicated double-mutants were treated with the indicated concentrations of each inhibitor for 4 h. Lysates were probed with the indicated antibodies. I) HER2 signaling and inhibitors of downstream pathways. J) MCF10A cells stably expressing HER2^L755S/T862A^ were treated with the indicated concentrations of each inhibitor for 3 days. Apoptosis was measured using the Caspase-Glo 3/7 assay and normalized to untreated cells. Data represent the mean ± SD (n=4). K) Cells were treated as in (J) for 7 days. Cell viability was measured using CellTiter-Glo. Data represent the mean ± SD (n=4). IC_50_ values (shown in parentheses) were calculated using GraphPad. ***p<0.001, ****p<0.0001, 2-way ANOVA + Bonferroni.

**Supplementary Figure 5:**
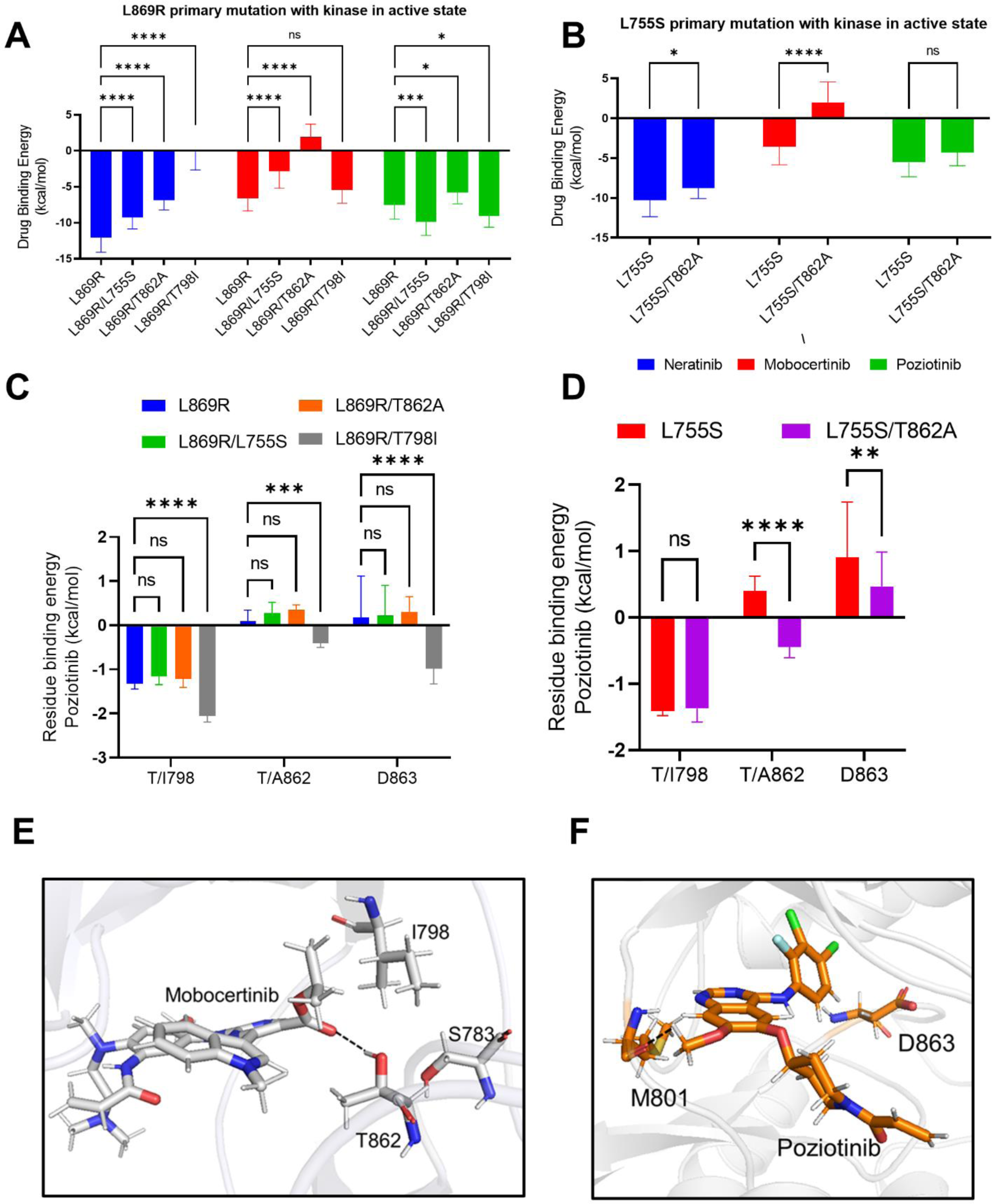
Acquired secondary HER2 mutations differentially impact binding affinity of HER2 TKIs. A) The Binding affinity of neratinib, mobocertinib, and poziotinib was calculated for HER2^L869R^ single- and double-mutants using MMPBSA. B) The Binding affinities for the HER2^L755S^ and HER2^L755S/T862A^ as in A). C-D) Energy decomposition analysis for HER2 residues T/I798, T/A862 and D863 for binding of poziotinib to HER2^L869R^ (C) or HER2^L755S^ (D) single and double mutants. The decomposition analysis was performed using MMPBSA. E) The structural model of mobocertinib for HER2^L869R/T798I^ to show the interaction of mobocertinib linker group with T862 and S783. The structure was obtained after 500 ns MD simulation F) The structural model of poziotinib for HER2^L869R/T798I^. The interaction between D863 and poziotinib is shown in the figure which seems to play important role for poziotinib binding with the gatekeeper mutation.

**Supplementary Figure 6.**
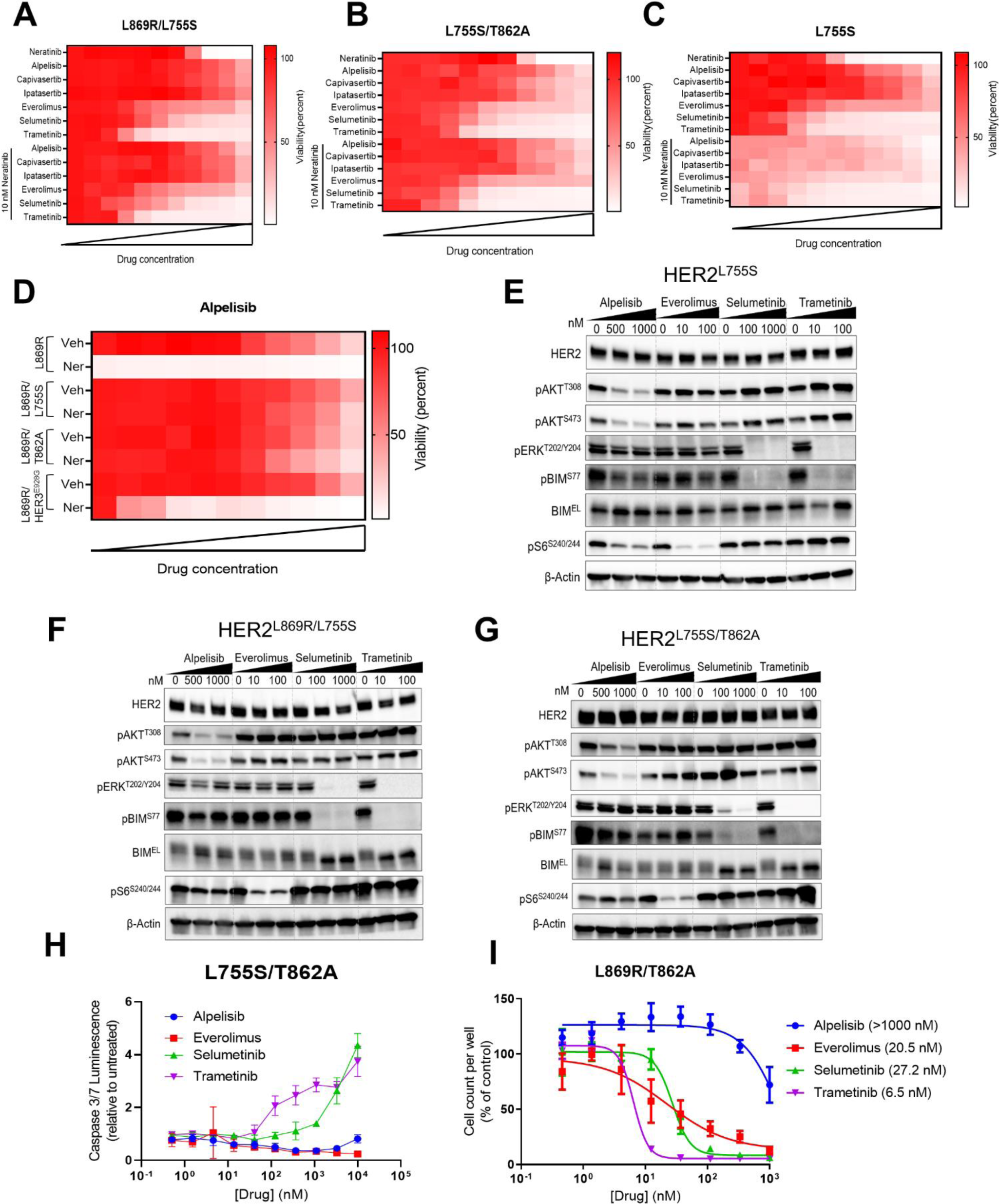
HER2 double mutants are sensitive to MEK Inhibitors. A-C) MCF10A cells stably expressing the indicated HER2 mutants were treated with increasing concentrations of each inhibitor (up to 10 μM) ± 10 nM neratinib in EGF/insulin-free media + 1% CSS for 6 days. Cell viability was measured by the CellTiter-Glo assay in quadruplicate. D) MCF10A cells stably expressing the indicated HER2 mutants were treated with increasing concentrations of alpelisib up to 10 μM ± 10 nM neratinib in EGF/insulin-free media + 1% CSS for 6 days. Cell viability was measured by the CellTiter-Glo assay in quadruplicate. E-G) MCF10A cells stably expressing HER2^L755S^ (E), HER2^L869R/L775S^ (F), or HER2^L755S/T862A^ (G) were treated with the indicated inhibitors for 4 h. Lysates were probed with the indicated antibodies. H) MCF10A HER2^L755S/T862A^ cells were treated with the indicated concentrations of each inhibitor for 3 days. Apoptosis was measured using the Caspase-Glo 3/7 assay and normalized to untreated cells. Data represent the mean ±SD (n=4). I) MCF10A cells stably expressing HER2^L869R/T862A^ were treated with the indicated concentrations of each inhibitor for 6 days. The number of cells/well was measured using Incucyte and normalized to untreated cells. Data represent the mean ± SD (n=4).

**Supplementary Figure 7.**
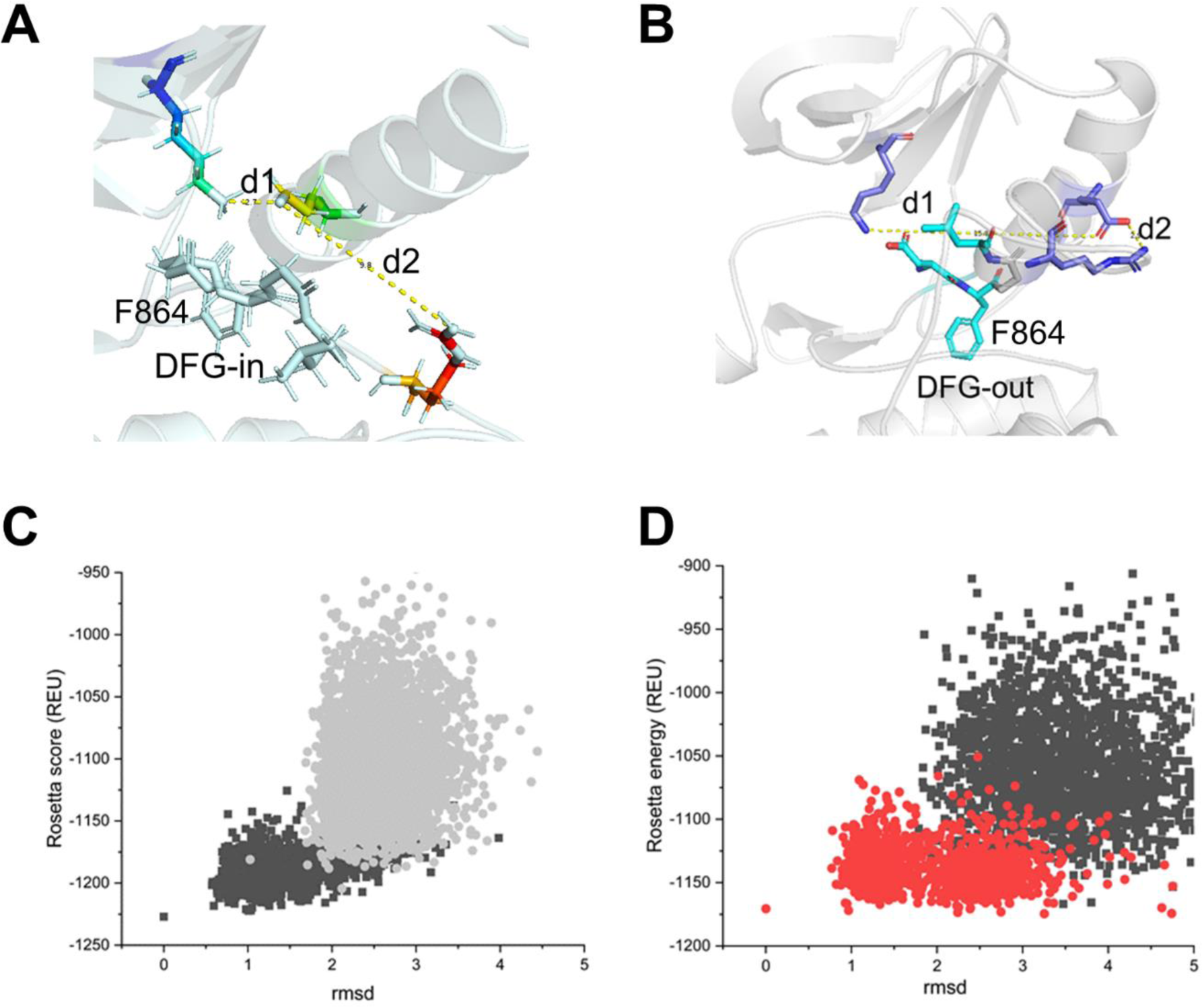
Computational methods. A) The active state model of HER2^WT^. The K753-E770 interaction is shown as d1 and the R868-E770 is shown as d2. The DFG-in motif is shown, which orients in an inward conformation in the case of active state B) The inactive state model of HER2^WT^. Residue F864 in the DFG-motif orients in the outward direction in the case of inactive state. C) The rmsd vs Rosetta score was obtained from the comparative modeling protocol in Rosetta. Two iterative round of CM modeling were performed to prepare the active state model of HER2, where the best scored models from the first round were used as seed for the second round of modeling. The 4riw and 3pp0 crystal structure was used as a template for CM. D) The inactive state comparative modeling for HER2, using a similar protocol as in (C). The CM modeling was performed using the 2gs7 structure as a template.

